# Shape, Shrink, Spheroid: A DIY High-throughput Spheroid Generation Device

**DOI:** 10.64898/2026.05.31.729042

**Authors:** Pankaj Mogha, Sourav Mukherjee, Tushar Gangwar, Debjyoti Roy, Aditi Vichare, Swanand Kulkarni, Vineet Sharma, Abhijit Majumder

**Author notes:** The authors contributed equally (Pankaj Mogha, Sourav Mukherjee, Tushar Gangwar).

## Abstract

3D spheroids, which closely replicate three-dimensional cell-cell and cell-extracellular matrix interactions, offer superior predictive capabilities compared to conventional 2D monolayer cultures, positioning them as forward-looking platforms in drug testing, cancer biology, and regenerative medicine. However, high-throughput generation of uniform sized spheroids is still a technological challenge. In one hand, the use of conventional ultra-low attachment (ULA) multiwell plates for this purpose is labour intensive and complex. On the other hand, the use of microfabricated facilities demands cutting-edge infrastructure such as clean room, photolithography, and microfluidic setup which are often unavailable for the resource constrained laboratories. In this study, we addressed these problems by developing a low-cost Do-It-Yourself (DIY), polydimethylsiloxane (PDMS) and agarose-based spheroid generation device, capable of producing and maintaining hundreds of spheroids with minimal user intervention. We have demonstrated two variants based on their size, termed here as S1 and S2 devices which fit into 6-well and 12-well plates, and can generate 600 and 1200 uniform-sized spheroids respectively. We validated our device with various cell lines including primary and cancerous cell lines. We further demonstrated the drug testing capabilities of the device by estimating the IC_50_ value of the anticancer drug Temozolomide on U87-MG. The value was comparable with the same obtained from the spheroids generated using conventional ULA plates. Additional attachment of a perfusion system made the device suitable for long-term spheroid culture without much user intervention. Furthermore, the devices can also be used for the production of spheroids with gradually changing diameters in a controlled manner, resembling a size gradient. This feature is useful for checking the effect of drugs on different-sized spheroids and for co-culturing spheroids with varying cell densities, mimicking the disease architecture. We have co-cultured two types of the placental trophoblast cells, i.e., extravillous trophoblast (HTR-8) and syncytiotrophoblast (BeWo) with varying densities. In summary, this paper demonstrates a unique DIY method for a high-throughput uniform-sized spheroid generation at a fraction of cost which can be deployed to resource-constrained labs.

## Introduction

Compared to the conventional 2D monolayer cell cultures, 3D spheroids preserve cell-cell and cell-extracellular matrix (ECM) interaction, mimicking the *in vivo* cellular microenvironment better [1]. The effects of hypoxia [2], inflammation [3], resistance to the drugs [2] [4], dynamic diffusion of cytokines [5] [6], etc., were shown to be more physiologically relevant in 3D spheroids compared to 2D monolayer cell culture. When cultured in 3D, senescence in the human mesenchymal stem cells was delayed and stemness was maintained for long-term [7]. Using glioblastoma as a model, it was demonstrated that aggressive characteristics of cancers are more prominent in 3D culture compared to their 2D counterparts due to the differential expression of long non-coding RNAs [8]. Similarly, in colorectal cancer, the dissimilarities in gene expression profile in 3D and 3D cultures have been shown by Abbas et al [9]. Due to these reasons, 3D spheroids are widely used to address several questions related to cancer biology [10] [11] [12], drug screening [13], cellular migration [14] [15] and secretome-based therapies [16] [5].

Over the past decade, several attempts have been made to generate 3D cellular models by integrating the knowledge of nanotechnology, bioengineering, material science and microfluidics. Commonly used techniques to generate spheroids include the hanging drop method [17], ultra-low attachment (ULA) plates [18], micropatterned plates [19], and liquid overlay methods [20]. In these methods, due to limited cell-substrate attachment, the cells come together to form 3D spherical aggregates or spheroids [21]. However, using these methods generating a large number of spheroids with minimal heterogeneity is laborious and time-consuming. Additionally, some of these methods have their own limitations such as culturing the spheroids for long-term with hanging-drop method becomes difficult and inefficient as the culture media in each droplet cannot be changed, resulting into increased osmolarity and cell death [22]. The rotating wall vessels are used for high-throughput and long-term culture of spheroids [23] but lacks spheroid compartmentalization and has size heterogeneity [24]. The microfluidic droplet-generation technique has been widely used [25] [26] for rapid generation of uniform size spheroids. In this technique, generally the cells are encapsulated inside droplets of hydrogel pre-polymer mix. Eventually, the droplets polymerize to provide 3D structures on which the cells grow into embedded spheroids. Recently, Namgung et al., have developed a DIY inexpensive setup, costing less than 7$ and capable of generating high-throughput spheroids through droplet formation by laminar jetting of cell suspension [25]. Many other microfluidic devices and spheroids-on-chip devices have also been developed in recent years, as reviewed in detail by Tevlek et al [27]. For example, Nguyen et al. [28], developed honeycomb shaped microwells using pre-stressed polystyrene sheets and laser-jet printer. Thomsen et al. [29], Marimuthu et al. [30] and Han et al. [31] produced high precision conical microwells array (CMA) using CNC controlled micro-milling machine, whereas Kim et al. [32] and Fang et al. [33] had reported fabrication of similar microarray for high-throughput spheroid generation by using resin and high precision 3D printer. In a recent study Chen et al. have used CAwell600 kit, a high-throughput microwell array for drug efficacy evaluation in liver steatosis and fibrosis [34]. Kim et al., have established spheroid co-culture model in a microfluidic chip consisting of colorectal cancer cells and primary rat liver cells to check the effect of drugs [35]. However, adopting these different fabrication techniques requires sophisticated instrumentation such as photolithography, cleanroom facility, CNC micromilling, high precision 3D printer, complex fluidic interfaces etc. making these methods inaccessible to resource limited laboratories and universities [36]. Furthermore, some of these fabrication techniques also have limitations related to portability, mass producibility, steep learning curve for the users, and difficulties in spheroid retrieval. Overall, these limitations hindered the mass adoption of such techniques for the research laboratories in general.

To address the above-mentioned infrastructure related challenges for the laboratories in LMICs, we have developed a simple Do-It-Yourself (DIY), low-cost, portable, microscopy friendly, microwell platform capable of generating high-throughput uniform-sized spheroids, eliminating the dependency on any sophisticated instrumentation such as photolithography, micromilling, or 3D printing. The novelty lies not only in its simplicity but also in its capability to deliver reproducible spheroids without the infrastructure overhead. For this purpose, the bottom side of a 384-well plate was used as the template on which Polyacrylamide (PAA) hydrogel was cast. Next, to miniaturize the structure, we adopted shrinkage-based microfabrication, as demonstrated by other groups [37], [38]. In our method, PAA due to its high degree of shrinkage [39] upon optimized controlled drying produced an array of microwells. The microwell structure thus obtained was used as mold to finally fabricate our device using soft lithography. By repeating the same replica molding-shrinking steps, it was possible to reduce the well size further. The devices such made are capable of generating hundreds of spheroids with minimal user intervention and in-device microscopic imaging including easy collection of the spheroids. We have demonstrated the capabilities with human mesenchymal stem cells (hMSCs), human umbilical vein endothelial cells (HUVEC), human glioblastoma multiforme cells (U87-MG), trophoblast cells (HTR-8, JAR and BeWo), mouse myoblast cells (C2C12), and human liver cancer cells (HepG2).

Additionally, a perfusion system was added for long-term culturing of spheroids with minimal user interference. We have also demonstrated that the device is capable of finding the IC_50_ of chemotherapeutic drug temozolomide and creating spheroids with size-gradients. Finally, we have created a 3D placental co-culture model with varying ratio of two types of trophoblast cells, extravillous trophoblasts (HTR-8) and syncytiotrophoblast (BeWo), mimicking the in-vivo interaction that occurs during the placentation process.

In summary, this paper demonstrates a novel and easy-to-fabricate method for high-throughput 3D spheroid culture, supporting both uniform and gradient size distribution, as well as the generation of co-culture models.

## Materials and Methods

### 1. Device Fabrication

The high-throughput spheroid generation device with microwells was fabricated by using a 384-well plate (Roche) as a template. To explain in brief, the patterns present on the back side of the 384-well plate was transferred to a polyacrylamide (PA) hydrogel which was then shrunk and used as a template for the next steps. PDMS: crosslinker (10:1 w/w) (Sylgard 184 from Dow Corning) was poured on the PA mold to produce a PDMS mold with 384 hills. This PDMS template (with hills) was then cut into desired shape (i.e., 10×10 for 100 wells) on which a second round of PDMS: crosslinker (10:1 w/w) was poured, producing the final S1 (size 1) device. Similarly, PA gel was poured on S1 PDMS mold (hills) and the downstream process was repeated to obtain the S2 (size 2) device which is smaller in size compared to S1 device. For more details, please refer to the ‘Result and Discussion’ section.

#### Fabrication of S1-sized agarose device

The process can further be simplified by using agarose as the final material instead of PDMS. Approximately 3–3.5 mL of the 2% agarose solution (HiMedia, MB002) in autoclaved ultrapure water was poured into each well of a 6-well plate. The PDMS stamp with microhills were securely placed onto these wells, making sure that it doesn’t completely touch the base of the plate. The plate was then transferred to a refrigerator set at 4°C for 5-10 minutes to complete polymerization. Following solidification, the PDMS mold was carefully removed, resulting into formation of microwells in the solidified agarose (supplementary Figure 1I). The agarose microwells were sterilized under UV light for an additional 30 minutes before using for spheroid culture.

### 2. Cell culture

The U87-MG cell line was a kind donation from Prof. Shilpee Dutt (JNU, India). High glucose Dulbecco’s modified Eagle medium (DMEM) (HiMedia-AL007A) was supplemented with 1% antibiotic-antimycotic solution (HiMedia-A002), 1% Glutamax (Gibco, 35050) and 10% foetal bovine serum (FBS) (HiMedia-RM1112) in a T25 flask for cell culture. Bone-marrow-derived hMSCs were purchased from Lonza (Cat. No. PT-2501) and were cultured in low-glucose DMEM (HiMedia, AL006) supplemented with 1% antibiotic-antimycotic solution (HiMedia, A002), 16% FBS (HiMedia, RM9955), 1% Glutamax (Gibco, 35050). JAR, HTR-8 and BeWo cells were generously provided by Dr. Deepak Modi (ICMR-NIRRCH, India). JAR, BeWo cells were cultured in DMEM/Nutrient Mixture F-12 Ham (DMEM/F12, 1:1 Mixture) (AL139A) supplemented with 10% FBS (HiMedia RM1112), 1% antibiotic-antimycotic solution (HiMedia-A002). HUVEC cells were cultured in HiMedia HiEndoXL Endothelial Cell Expansion Medium (AL517) supplemented with 1% antibiotic-antimycotic solution (HiMedia-A002), All the cells were cultured under humidified conditions at 37°C with 5% CO_2_.

Once confluent, cells were trypsinized using TrypLE™ (Gibco, 12604021) and was neutralized with complete media of the respective cell types. For estimating the cell count, the cell suspension was centrifuged at 1200 rpm for 5 minutes and was counted using a hemocytometer (Invitrogen) for further studies.

### 3. Generation of 3D spheroids

PDMS devices were treated with air plasma (Harrick Plasma) for 2 minutes making the surface activated and hydrophilic. Immediately after plasma exposure, the device was incubated in 70% ethanol for 20 minutes for sterilization. The device was rinsed thoroughly with DPBS (Dulbecco’s phosphate buffered saline) (HiMedia-TS1006) at least three times to remove any ethanol residues. The device was then filled with a 2% Pluronic F-127 (Sigma-Aldrich) solution in Milli-Q water and incubated at 37°C for 2–3 h to avoid any cell attachment at the bottom of the microwells. Post incubation, the device was washed with complete media at least three times to remove any remaining Pluronic.

Next, in 2 mL and 1 mL of media with 0.1 × 10^6^ cells were seeded for 100 well S1 and S2 devices respectively. To achieve the spheroid size gradient, 0.55 × 10^6^ cells in 2 mL media were seeded in a S1 agarose device (110 wells). For optimal cell viability, 1 mL of the media was replaced daily.

### 4. Addition of flow system

To change the media continuously, the device assembly was connected with a syringe pump (New Era Pump Systems Inc., Model NE-1000). Media inlet and outlet tubing were connected with the device by punching two holes on the opposite sides of the device. The height of the outlet was lower than the inlet height. This allowed the continuous media flow without requiring a lid or making it a closed device. The flow was turned on at a flow rate of 84 µL/h post 24 hours of cell seeding and maintained for the whole duration of spheroid culture. The whole setup was placed in a 150mm petri-dish to maintain sterility for long-term spheroid culture.

### 5. Generation of gradient-sized spheroids

The S1 agarose devices were inclined during cell seeding to achieve spheroids of size gradient. HTR-8 cells were trypsinized and 0.55 × 10^6^ cells were seeded in S1 agarose device with 110 wells. Agarose devices were preincubated with complete media to prevent bubble formation while cell seeding. Cells were then seeded in the wells from the bottom portion of the device while maintaining the incline of 20°. After 12 h of incubation, the devices were transferred to a flat surface, and 1 mL of the media was replenished daily to maintain cell viability.

### 6. Co-Culturing of Spheroids in the S1 Device

To generate 3D co-culture spheroids, extravillous trophoblast cells (HTR-8/SVneo) and syncytiotrophoblast-like BeWo cells were differentially labelled using CellTracker™ dyes. HTR-8 cells designated as the spheroid core, were labelled with green cell tracker and the BeWo cells forming the outer layer were labelled with a red cell tracker. The HTR-8 cells were trypsinized and pelleted down. The cell pellet is then resuspended and incubated in a pre-warmed CellTracker™ Green CMFDA solution (10 μM; Invitrogen, C2925) for 30 minutes at 37 °C. The staining reagent was prepared in complete culture medium. Post-incubation, cells were washed with fresh medium, centrifuged at 1200 rpm for 5 minutes, and resuspended for seeding. Around 6 × 10^5^ green fluorescence labelled

HTR-8 cells (per device of 120 wells) were seeded into the S1 spheroid culture device in either a gradient or flat distribution pattern, allowing control over core cell density. Spheroids were allowed to self-assemble and compact over a period of 4 days. Subsequently, BeWo cells were stained with CellTracker™ Red CMTPX (10 μM; Invitrogen, C34552) using the same protocol as above and introduced into the device to initiate co-culture. In the first setup, BeWo cells were seeded flat onto the pre-formed HTR-8 gradient cores. Whereas, in the second setup HTR-8 cells were seeded flat, and BeWo cells were introduced using a gradient seeding pattern. Following seeding, the co-cultures were incubated for an additional 4 days to promote spheroid maturation and cell-cell interaction. The resulting dual-colour spheroids were imaged using a confocal laser scanning microscope with z-stack acquisition. Fluorescent signals were captured using excitation/emission filters suitable for green (approximately 492/517 nm) and red (approximately 577/602 nm) fluorophores.

### 7. Spheroid harvesting

Once the spheroids were formed within the device, the existing culture medium was removed by tilting the device. Subsequently, fresh culture medium was forcefully introduced into the device using a 1 mL pipette, which effectively dislodged the spheroids from the microwells, enabling their collection in a separate Petri plate. If required, individual spheroids could be isolated by aspirating them directly with a 200 µL pipette.

### 8. Drug Testing with cultured spheroids

To evaluate the cytotoxicity of Temozolomide (TMZ) (CombiBlocks, Cat. No. OR-2567) on U87-MG cells, drug stock solutions were prepared in DMSO (Sisco Research Laboratories, Cat. No. 30239). U87-MG cells were seeded in each well of a 96 well for 2D-monolayer formation. For 3D spheroid generation, 1000 cells were seeded in each well of a ULA plate and 0.1 × 10^6^ cells were seeded in S2 devices. The spheroids were cultured for 7 days in complete DMEM media, replenishing the media daily. TMZ was tested across concentrations from 0 to 800 μM. Media and vehicle control wells were included, and DMSO levels were controlled to not exceed 1% per well. Cells were incubated with the drug for 48 h in the incubator at 37°C with 5% CO2. After 7-days of spheroid culture, the spheroids were taken out of the S2 device and kept in a 96-well plate. Following the 48-h of TMZ incubation, the media was removed, and 100 μL of freshly prepared, 5 mg/mL MTT solution (3-(4,5-dimethylthiazol-2-yl)-2,5-diphenyl tetrazolium bromide, Sigma Chemical Co., St Louis, MO, USA) in DPBS was added. The plates were incubated for 1–2 h at 37°C to allow formazan crystals to form. After incubation, the MTT solution was removed, and 100 μL of DMSO was added to each well to dissolve the crystals. Absorbance was measured at 570 nm using a spectrophotometer. These absorbance values were used to assess cell viability, quantifying drug-induced cytotoxicity. All steps involving DMSO were conducted under reduced light exposure to minimize compound degradation. The IC_50_ values were calculated by fitting the percentage of viability in three parameter model by using “Quest Graph™ IC_50_ Calculator” (AAT Bioquest, Inc., https://www.aatbio.com/tools/ic50-calculator). Following is the formula used for fitting and identifying the IC_50_.

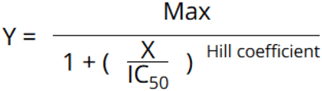

### 9. Spheroid staining and image analysis

For Hoechst 33342 (Life Technologies, Thermo Fisher Scientific) staining (1:5000), Calcein-AM (Life Technologies, Thermo Fisher Scientific) staining (1:2000), and Propidium Iodide (PI) (Life Technologies, Thermo Fisher Scientific) staining (1:1000) cells were incubated for 30 minutes and fluorescent images were captured with EVOS FL Auto fluorescence microscope (Life Technologies, Thermo Fisher Scientific), while phase contrast images were captured with Nikon Eclipse Ti2 microscope. For the characterization of the spheroids, the area of the spheroids was measured by tracing the boundary of the spheroids with ‘Freehand selections’ tool in ImageJ (National Institutes of Health, Bethesda, USA). The diameter of the spheroids was calculated from the obtained projected area. The following equation was used for calculating the diameter.

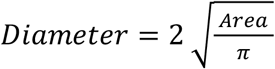

## Results and Discussion

### 1. Fabrication of the device

To fabricate the high-throughput spheroid generation device, the bottom side of the commonly available 384-well plate was used as the template which has 384 raised hill-like structures arranged in a 16 × 24 matrix. As the apex of those hill-like structures were flat, to obtain the round bottomed well, the flat hills were coated with small drops of nail paint (Figure 1A-B, Supplementary Figure 1A (i) and B). Once the nail paint dried, polyacrylamide (PA) gel solution, consisting of acrylamide (8%), bis-acrylamide (0.48%), APS (1:100), and TEMED (1:1000), was poured onto the backside of the well plate and allowed to solidify (Figure 1C-D). This particular gel composition gives rise to a PA gel of rigidity 40 kPa when solidified [40] which was found to be optimal for faithful transfer of the structures, material strength, and sufficient shrinking. We tried the gel formation with stiffness lower than 40 kPa such as 20 kPa and 5 kPa. The PA gels with lower stiffness were brittle and tough to handle for downstream processes (Supplementary Figure 1G). Hence, further experiments were done with 40 kPa gels. After solidification, the hydrogel was carefully removed and submerged in absolute ethanol to induce dehydration-mediated shrinkage. Ethanol, being a non-aqueous, hydrophilic solvent, replaces the water within the hydrogel network due to its higher affinity and miscibility with water. This exchange leads to the collapse of the polymer network and isotropic shrinkage of the gel. To minimize the formation of stagnant boundary layers around the gel surface and ensure uniform solvent exchange throughout the gel, the system was placed on a rocker, providing gentle agitation. Fresh ethanol was introduced at least three times at 3–4 h intervals to maintain a sufficient chemical gradient for effective dehydration and to prevent local saturation effects, thus promoting homogeneous shrinkage across the entire gel volume (Figure 1E-F). This step was crucial as not changing the ethanol leads to incomplete dehydration and shrinkage of the PA gel (Supplementary Figure 1H). Overall, the ethanol assisted shrinkage eliminated the distortion or crack formation commonly observed with air drying (Supplementary Figure 1E).

**Figure 1:**
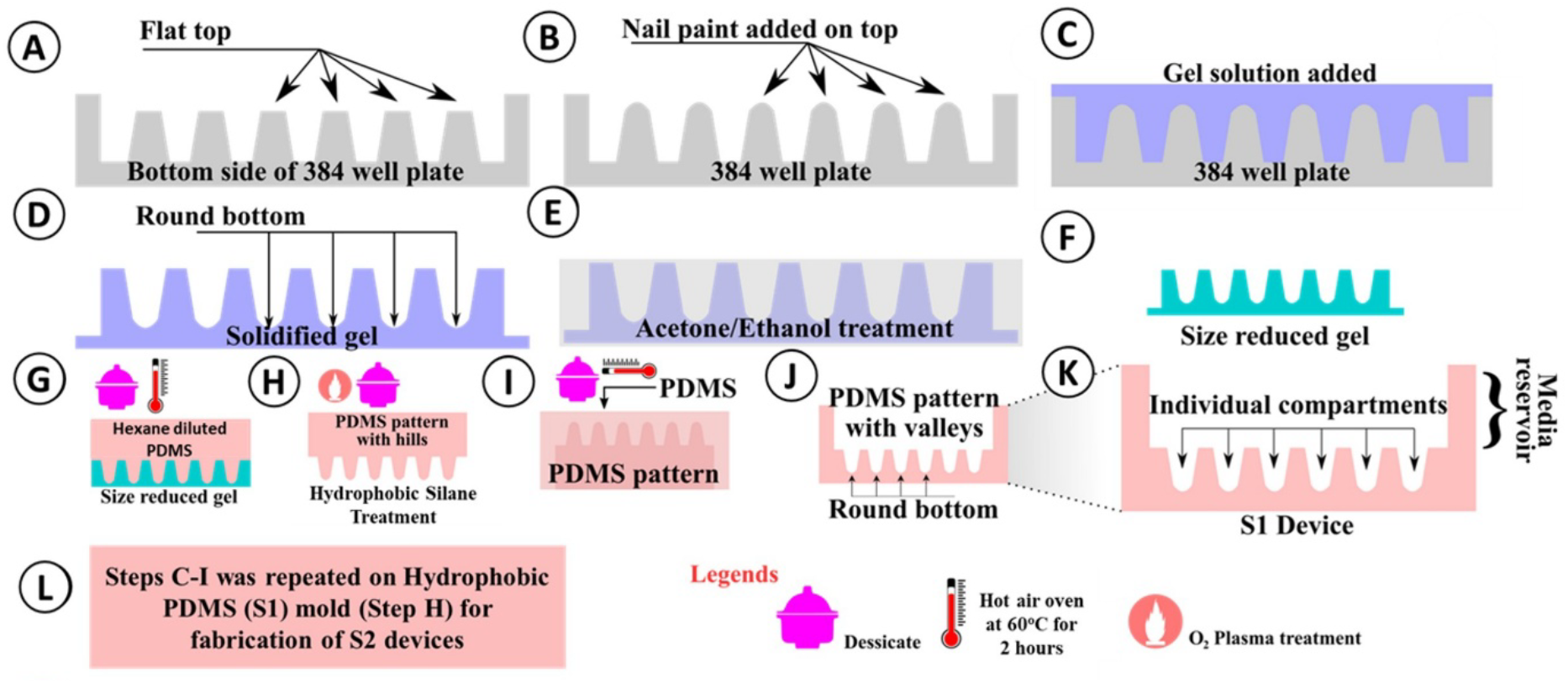
Steps of the device fabrication: Schematics showing (A) the cross-section of an inverted 384 well plate (bottom side up). (B) Nail paint coated rounded top. (C) Polyacrylamide gel solution poured on the well plate. (D) Upon solidification, the gel structure with round bottom wells were achieved. (E) The gel was incubated in ethanol overnight with replacement of ethanol at least three times for drying and shrinking. (F) The size-reduced gel was then removed from ethanol and air dried. (G) Hexane-diluted PDMS was then poured onto the shrunk gel (mold), degassed, and cured at 60°C overnight. (H) After curing the PDMS was removed from the mold and turned hydrophobic by incubating in silane under vacuum for 4 h after a brief exposure to air plasma. (I) The hydrophobic PDMS was then used as mold and PDMS was poured, degassed, and cured at 60°C overnight. (J) The cured PDMS was then removed carefully from the hydrophobic PDMS which resulted in the fabrication of a PDMS device with 384 round bottom wells. (K) Enlarged schematic of the 384 well device where the compartment above the microwells is used as a media reservoir. (L) Step C to I were repeated with the S1-device as a mold to get a 2° shrunk device, called S2-device (Size 2).

Once the gel had undergone ethanol-induced shrinkage, it was further dried in a hot-air oven at 60°C to remove residual bound ethanol from the polymer network. This combined dehydration process transformed the initially soft, water-rich polyacrylamide (PAA) hydrogel into a hardened and mechanically stiff mold. To correct any curvature or warping that developed during the shrinkage process (Supplementary Figure 1F), the dried PAA structure was sandwiched between two tightly fastened glass slides and placed in a closed humid chamber. When this assembly was kept in a hot-air oven for 3–4 hours, the humid environment partially rehydrated the polymer matrix, temporarily plasticizing it and increasing its flexibility, thereby allowing uniform straightening under mechanical constraint. Following this relaxation and shape-correction step, the PAA mold was subjected to a final drying cycle in the hot-air oven at 60°C for 2–4 hours, yielding the stable, miniaturized, and planar final structure (Supplementary Figure 1A(ii)). Next, PDMS base and crosslinker were mixed at a 10:1 weight ratio (monomer:crosslinker) and degassed under vacuum to eliminate air bubbles introduced during mixing. To reduce the viscosity and enhance the ability of the PDMS mixture to infiltrate the fine features of the PAA mold, the degassed PDMS was further diluted with n-hexane at a 10:3 volume ratio (PDMS mixture: hexane). It was observed that 10:3 ratio provided the optimum clarity, easy removal of trapped air, and faithful pattern transfer, unlike the other combinations and undiluted PDMS (Figure 1G, Supplementary Figure 1C, J). The diluted PDMS was then poured onto the PAA mold in successive layers, with intermittent degassing steps to ensure the complete removal of trapped air, particularly from the bottom of the microwells (Figure 1G, Supplementary Figure 1C). Following degassing, the assembly was placed in a hot-air oven at 60°C and cured overnight, allowing gradual crosslinking of the PDMS while permitting slow evaporation of hexane, thus minimizing the formation of voids or defects in the final elastomeric structure. The cured PDMS (with hills) was then carefully removed (Supplementary Figure 1A(iii)), cut into smaller pieces as per requirement such as 10 × 10 (100 microwells) devices and used as mold for further experiments (Supplementary Figure 1A (iv)). Next, these PDMS hills (mold) were made hydrophobic by submerging them in trichloro (1H,1H,2H,2H-perfluorooctyl) silane diluted in hexadecane (ratio of 1:100 v/v) for 2 h at room temperature (Figure 1H). This silane treatment is crucial to make the PDMS mold (with hills) hydrophobic, negating the PDMS-PDMS attachment while separating the device from the mold. Post-treatment, the mold was incubated for 24 h at 60°C. This PDMS mold was then used as a template for generating the final S1 (size 1) device as mentioned. The PDMS: crosslinker (10:1 w/w) was poured onto the hydrophobic PDMS mold, degassed, and cured at 60°C overnight (Figure 1I) to generate the first order shrinkage device (S1 device) with microwells (Figure 1J-K, Supplementary Figure 1A (v)). It is to be noted that this hydrophobic PDMS mold can be reused multiple times.

To fabricate the second-order shrinkage device, referred to as the S2 (size 2) device (Supplementary Figure 1A(viii)), the hydrophobic S1 PDMS mold with raised features was used as a template. Notably, unlike the S1 fabrication steps, the use of the original 40 kPa polyacrylamide (PAA) gel formulation was found inadequate for replicating the finer structural details required in the S2 device (Supplementary Figure 1K). To address this, the water content in the gel formulation was reduced to increase stiffness, and a higher-modulus PAA hydrogel was prepared by mixing 40% (w/v) acrylamide and 2% (w/v) bis-acrylamide in a 1:1 ratio. Polymerization was initiated by adding 10% ammonium persulfate (APS) at 1:100 (v/v) and tetramethylethylenediamine (TEMED) at 1:1000 (v/v). This formulation, corresponding to an approximate elastic modulus of ∼100 kPa, was critical for achieving high-fidelity pattern transfer from the PDMS template to the hydrogel. Following gel polymerization, steps E–I (as described above) were repeated to generate the second-order shrinkage device (S2 device) (Figure 1L and Supplementary Figure 1A (vi–viii)).

While the shrinkage-based microfabrication has been demonstrated by other groups [37], [38], they have not utilized for spheroid generation. To be used as microwells which may give rise to uniform sized spheroids, the devices need to be perfectly un-distorted, defect free, the surface walls need to be crack free, and smooth. We achieved these objectives, by careful optimization of the drying, shrinking, and pattern transfer process as explained above.

### 2. Characterization of the device

To demonstrate the size of the devices, it is compared with a one-rupee coin (INR) having a diameter of 2 cm (Figure 2A). The length and width of the 10 × 10 S1 and S2 PDMS devices, each with 100 microwells, were measured to be 2 cm x 2 cm and 0.8 cm x 0.8 cm, respectively (Figure 2B-C). A 10 × 10 (100 microwells) S1 device fits in a 6-well plate, while the S2 device fits in a 12-well plate (Figure 2D-E). The device dimensions are not limited to the mentioned sizes but can be adjusted to any user-defined A x B size (Supplementary Figure 1A(iv)). After the first shrinkage, the area of the 384-well plate, initially 94 cm^2^, reduced to approximately 20 cm^2^, which is 79% shrinkage. Similarly, after the second round of shrinking, the area reduced from 20 cm^2^ to 6.6 cm^2^ which is approximately 70% shrinkage, shown in Figure 2F-H. The volume of the S1 and S2 microwells were measured to be approximately 5µL and 1µL respectively. This shrinking allows us to seed the cell suspension at once unlike 96-well ULA plates where one needs to seed in 100 wells separately to obtain 100 spheroids. Due to shrinkage, the inter-well distance is reduced, allowing the cells to get distributed and settled in 100 wells to form 100 spheroids in one device placed in one well of 6-well plate. Thus, not only reduces the space consumption from six 96 well plates to one 6-well plate but also saving time and labor for the users. To characterize the well dimension, the S1 and S2 devices were longitudinally cross-sectioned and imaged under a microscope. The imaging confirmed the conical shape of the wells with round bottom (Figure 2I). The bottom diameter of S1 and S2 microwells were approximately 800 µm and 500 µm respectively (Figure 2J-K). The top diameter of the S1 and S2 microwells were approximately 1500 µm and 850 µm respectively. The height from the bottom to the top of the well was found to be approximately 3 mm and 1.2 mm respectively. To compare the well and spheroid size, a schematic of S1 and S2 microwells is shown in Figure 2L containing spheroids with an average size of 300 µm.

**Figure 2:**
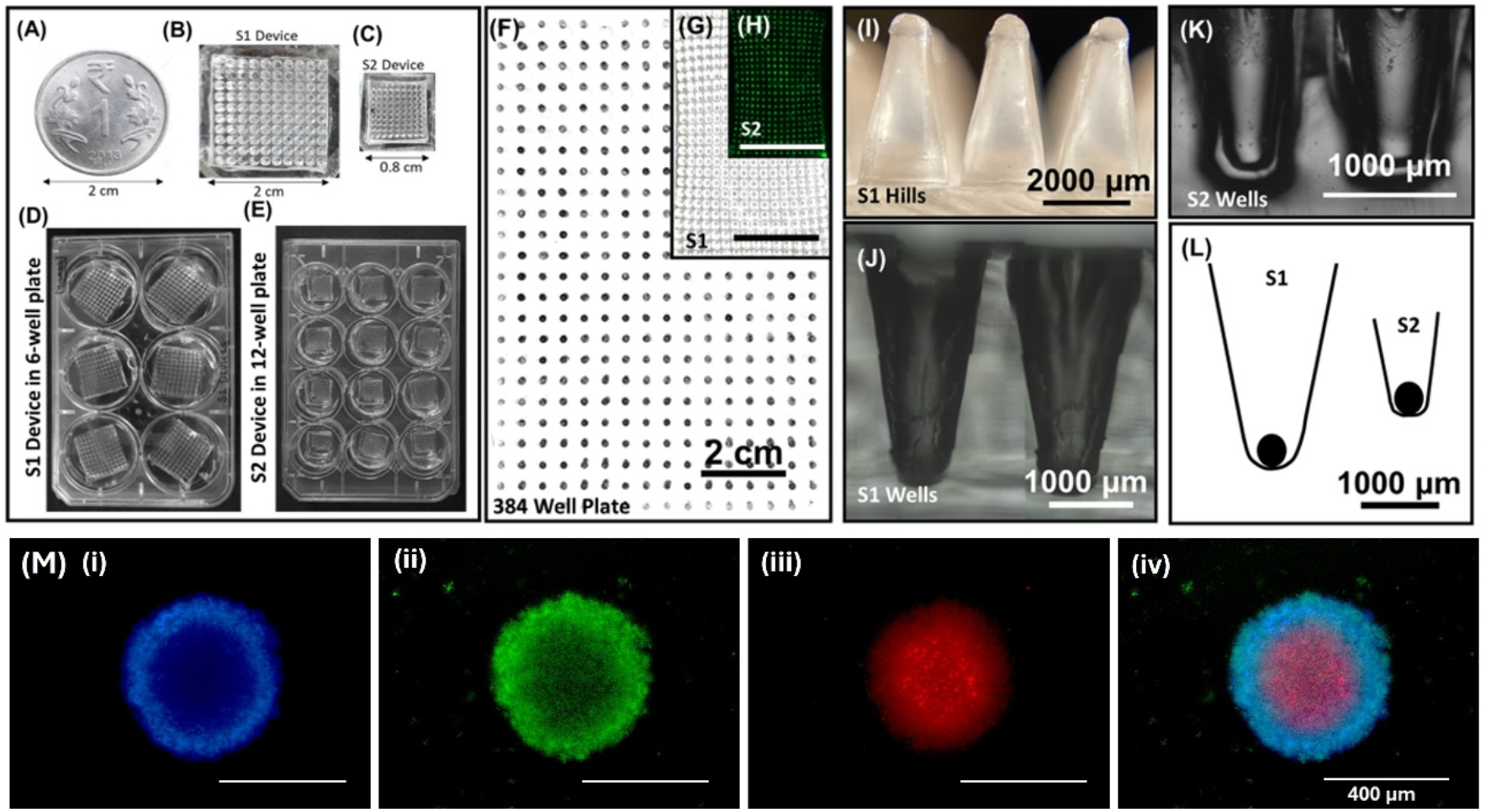
Dimensions of fabricated device and microwells: (A) A one rupee coin (INR), (B) Real image of 10 × 10, 100 well S1 device, (C) Real image of 10 × 10, 100 well S2 device. (D) Six S1 devices fitted into a regular 6-well plate, (E) Twelve S2 devices fitted into a regular 12-well plate, (F) The bottom side of the 384 well plate. (G) Shrunk PAA gel (S1) after treatment with ethanol. (H) Fluorescent image (green) of the PDMS-S2-device, each well containing FITC solution. The scale bar for F-H is 2 cm. (I) Image of hills of PDMS-S1 mold. Scale bar 2000 µm. (J and K) The cross-sectional image of S1 (J) and S2 (K) device microwells. Scale bar 1000 µm. (L) Representative drawing of the S1 and S2 device microwells occupied with spheroids, (M) Fluorescent labelled 2Dimages of U87-MG spheroids. The day 7 spheroids were stained with (i) Hoechst (blue), (ii) Calcein-AM (green), and (iii) Propidium iodide - PI (red), (iv) the merged image of all three channels has been shown, scale bar 400 µm.

To demonstrate the universality of devices, spheroids of different cell types were formed. One such typical spheroid from U87-MG glioblastoma cells stained with Hoechst, Calcein-AM and PI after 7 days of culture is shown here (Figure 2M). The spheroids from tumor cells, or often called tumoroids, are known for the three layers of cells. The uppermost layer is made of proliferative cells, followed by quiescent layer and a necrotic core. Cells in both proliferative and quiescent layers are viable. However, cells at the core are dead or dying due to the non-availability of oxygen and nutrients [41] which is a typical signature of *in vivo* tumor [41] [42]. The U87-MG spheroids formed using the current device show the necrotic core (in Red, PI) and viable outer layers (in Green, Calcein) establishing the physiological relevance of the model (Figure 2M). Furthermore, the spheroids were imaged in the PDMS device under a fluorescent microscope, demonstrating the capability of in-device imaging. As the device is made of PDMS, the bottom of the wells is transparent. This transparent well allows full device imaging (Supplementary Figure 2B) of the spheroids which can be extremely useful to assess the effect of drugs on cytotoxicity, or to check the spheroid growth over days. As the devices are kept in regular multiwell plates, the whole system is portable, and ready for live cell imaging or time lapse imaging using any compatible microscope. Such real time imaging is not possible in rotating wall vessels or might be quite difficult and demands customization in complex microfluidic structures [27].

### 3. Spheroid Formation and their Characterization

Two major areas of 3D spheroid application are tumoroid and organoid generation. Hence, the universality of the device was demonstrated by utilizing multiple cancer cell lines such as glioblastoma multiforme U87-MG, choriocarcinoma BeWo and JAR (Supplementary Figure 2-A), and with bone-marrow derived human mesenchymal stem cells (hMSCs). Other than cancer cell lines and stem cells, spheroids of HUVEC cells were also formed using this device. However, for brevity, data with U87-MG and hMSC spheroids are presented in this paper.

To form spheroids, two conditions are to be satisfied. First, the cells should come together, which we achieved by creating U-shaped round bottomed, conical microwells, as shown in Figure 2J-K. The second condition is that the cell-cell adhesion should be more than cell-substrate adhesion. To fulfill this criterion, we made the devices non-adhesive by treating them with air plasma followed by 5% Pluronic treatment for 20 minutes at room temperature (see method for more details and Supplementary Figure 1-D). After this step, cells were seeded at a density of 1000 cells/well i.e., 0.1 × 10^6^ cells in a device with 100 wells (Figure 3A). Spheroid with lower seeding densities (500 cells/well and 100 cells/well) were also easily achieved (Supplementary Figure 2C).

**Figure 3:**
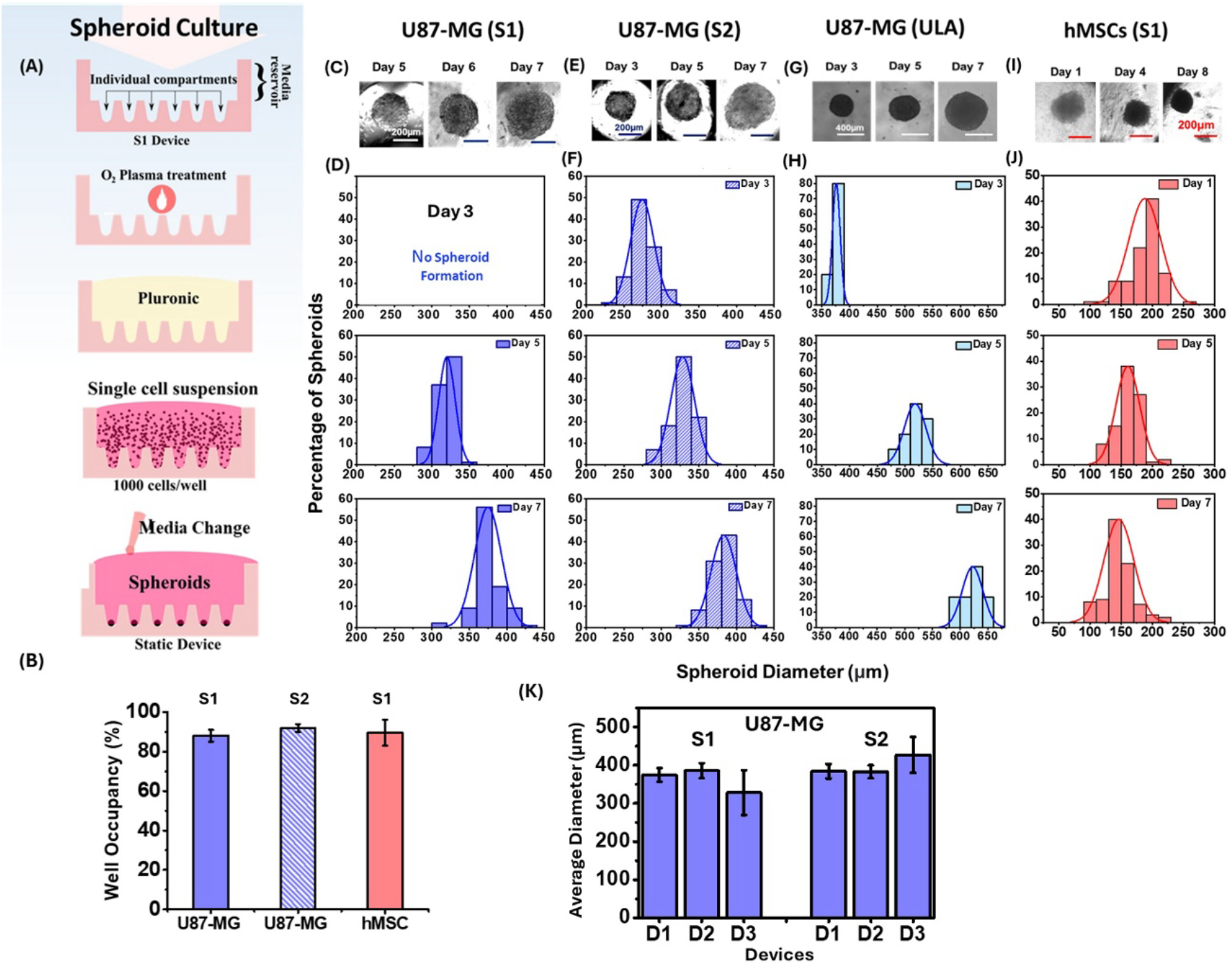
Spheroid distribution, well occupancy and size distribution: **(A)** Schematic for spheroid formation and culture showing the individual compartments of the device, air plasma, Pluronic treatment, cell suspension after seeding, spheroid formation, and media change. **(B)** The percentage of well occupancy of S1, S2 devices with U87-MG (blue bar) and hMSC spheroids (red bar). Data represents mean ± standard deviation (SD) and n=5. **(C)** Phase contrast images and **(D)** size distribution of U-87 MG spheroids in S1 device for different days of culture, scale bar is 200 µm. Similarly, phase contrast images and size distribution of U87-MG spheroids in the S2 device (blue-patterned), ULA plate (cyan) (n=6), and hMSCs in S2 devices are shown in **(E-F), (G-H)**, and **(I-J)** respectively (n>90). For ULA plate (G), the scale bar is 400 µm. **(K)** Device to device variation and the effect of user handling of S1 and S2 are represented by the mean spheroid diameter of U87-MG spheroids on day 7 in three devices (D1, D2 and D3). Data represents mean ± standard deviation (SD) from (B) devices (n>3) and (K) spheroids (n>90).

To evaluate the effectiveness of Pluronic coating in preventing cell adhesion and ensuring uniform spheroid formation, various surface treatment conditions were tested: UV exposure (30 minutes) followed by coating with 2%, 5%, or 10% Pluronic, and plasma treatment (2 minutes) followed by coating with 2% or 5% Pluronic. The treated devices were incubated at 37°C for 2 hours. Among these, the condition involving 2-minute plasma treatment followed by 5% Pluronic coating resulted in minimal cell attachment and consistent, intact spheroid formation. In contrast, other conditions showed noticeable cell adhesion to the well surfaces, leading to irregular or multiple small spheroids within individual wells. (Supplementary Figure 1-D)

We benchmarked our device against conventional 96-well ultra-low attachment (ULA) plates. Unlike ULA plates, which require the separate addition of cells into each well, our devices—owing to their compact size—accommodate 100 microwells within a single well of 6-well or 12-well plates (for the S1 and S2 devices, respectively). This design enables simultaneous addition of cells and media to all 100 microwells. The obvious question arises whether all the microwells would receive the cells and would be able to form a spheroid. We observed that 88 ± 3% and 92 ± 2% microwells were occupied with U87-MG spheroids in S1 and S2 devices respectively. Similarly, the hMSC spheroids in the S1 devices showed 87 ± 4% well occupancy (Figure 3B). Only a few wells at the edges did not receive the cells or failed to form a spheroid.

Next, to check the spheroid growth U87-MG cells and hMSCs were cultured in the device for one week and phase contrast images were captured every day. Both S1 and S2 devices as well as ULA plates showed a steady increase in the spheroid size over days indicating cell proliferation (Figure 3C). The average diameter of the U87-MG spheroids in the S1 device increased from 320 ± 11 µm at day 5 to 374 ± 18 µm at day 7 (p-value <0.001) (Figure 3D). Similarly, in the S2 devices spheroid diameter increased from 274 ± 15 µm on day 3 to 382 ± 17 µm on day 7 (p-value <0.001) (Figure 3E-F), and in the ULA plates it increased from 376 ± 6 µm on day 3 to 622 ± 18 µm on day 7 (p-value <0.001) (Figure 3G-H). Interestingly, although the microwells were of different sizes, both S1 and S2 could produce spheroids of similar sizes which were smaller than ULA formed spheroids. While both S1 and S2 devices produced spheroids of similar sizes, the kinetics were different. We observed that the U87-MG cells started forming spheroids on day 5 in S1 devices while in S2 devices and ULA plates it started on day 3 (Figure 3C-H). Due to their smaller size, S2 could bring the same number of cells together faster compared to the S1 device. However, S1 devices can also produce spheroids within day 3 if cell seeding density is increased (data not shown). Although the S2 wells are smaller in size compared to S1, the spheroid-forming ability of U87-MG cells remained unchanged. As a result, both devices produced spheroids of similar size by day 7. It is to be noted that, S1 can accommodate larger spheroids comfortably, while S2 supports relatively smaller spheroids due to smaller well size.

To demonstrate the universality of devices at generating spheroids primary cells were used, similar experiments were performed using human mesenchymal stem cells (hMSCs). Unlike U87-MG, hMSCs started forming cell aggregates on day 1 in S1 devices (Figure 3I). The diameter of the hMSC spheroids exhibited wider distribution on early days (day 1) which progressively narrowed with passing days, where spheroid size distribution was more uniform on day 8 (Figure 3J). Interestingly, the size of the spheroids reduced with days as the spheroids became more compact which is evident from the change in spheroid transparency. The average diameter was reduced from 189 ± 31 µm on day 1 to 146 ± 23 µm on day 8 (p-value <0.001) (Figure 3J). This reduction in the diameter of the spheroids formed with primary cells has been previously reported by Rovere et al. [43] and Deynoux et al [44].

Device-to-device and user-to-user variability, two critical parameters for any fabrication process were estimated (Figure 3K and Supplementary Figure 2D). Here, the first two devices were fabricated and used by user1 (well-trained) whereas the 3rd device was fabricated and used by user2 (untrained). No major dependency of mean spheroid diameter on the batch of manufacture or training level of the user was observed demonstrating that these spheroid devices can be used by individuals familiar with basic cell culture techniques but without any device-specific training. Furthermore, we shipped the devices (or trained to fabricate the device) to our collaborators Dr. Deepak Modi (NIRRCH-ICMR, Mumbai) and Prof. Sudhir H Ranganath (Siddaganga Institute of Technology, Tumkur, India) for evaluating the devices independently. These independent laboratories also could use our method/device for their 3D culture of placental trophoblasts and glioblastoma cells respectively (data not shown for confidentiality).

Another important feature to observe from this data is spheroid homogeneity. On day 7, for the trained user, the standard deviations for S1 and 2 devices were within 5% of the mean. For a conventional ULA plate, the same was ∼4%, establishing the usefulness of the device to create monodispersed spheroids. However, for the untrained user, the standard deviation was larger (18% for S1 and 11% for S2) compared to that of the trained user indicating that the user preparedness might have some effect on spheroid homogeneity. To note, for the same footprint as a 96 well ULA plate, S1 and S2 devices can produce 600 and 1200 spheroids respectively, highlighting the high throughput capability of the devices. Additionally, all the spheroids can be collected by simply tilting the device, or from each microwell.

### 4. Device with continuous perfusion

3D spheroids are often cultured for long durations which demand regular replenishment of the media [45]. Also, for drug testing or cytotoxicity testing applications, media is to be changed from growth media to media with specific drugs. For organoid cultures, changing media from growth to differentiation media might also be required [46]. For more complex treatments/analyses, one may need to change to different media compositions at different time points [47] [48]. All such liquid handling is tedious, demands a significant amount of person hours, and may cause the loss of spheroids in a conventional ULA plate. To address these issues, a perfusion system was added to the S1 device for the automated replenishment of culture media. The perfusion system, connected to a microfluidic syringe pump, was turned on post 24 hours of cell seeding. This time gap allows the cell suspension to settle at the bottom of the wells. Once the pump was turned on, no further user attention was provided to the device. U87-MG and hMSCs were cultured for 8 days in the S1 device with continuous perfusion at the flow rate of 84 µl/h which allowed 2 mL of media replenishment every day (Figure 4A-i-iii), equivalent to complete media change in every 24 hours. Simultaneously, static devices were also maintained alongside devices where media was not changed for 2 and 4 days consecutively. By positioning the outlet at a lower height than the inlet, continuous perfusion of the culture media was maintained, even though the device remained open.

**Figure 4:**
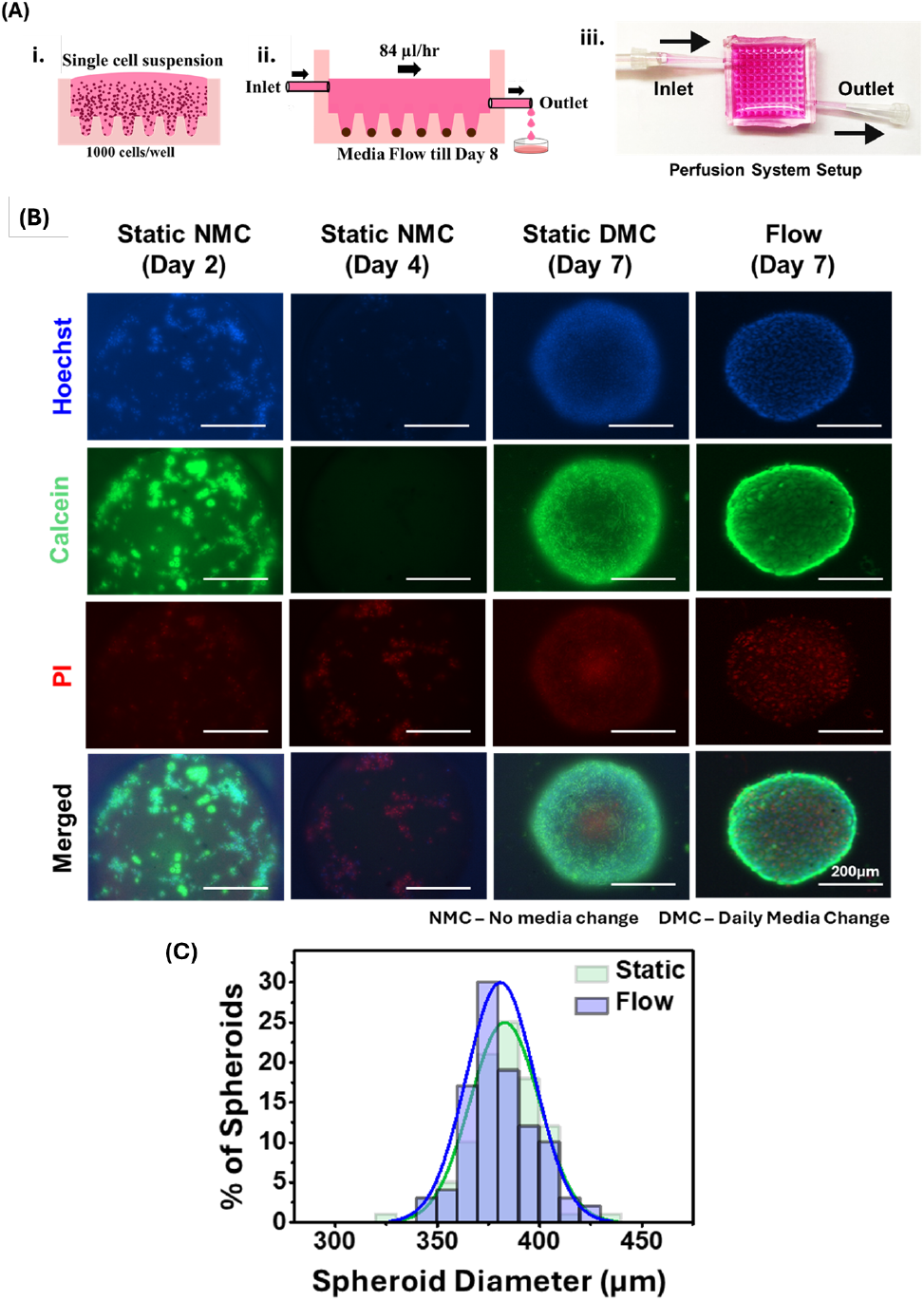
Long-term spheroid culture with perfusion system: **(A) i. and ii** show the schematic of spheroid culture in the flow device. The outlet are placed at a lower position to allow continuous flow in an open device setup, **iii**. An actual image of the flow setup. **(B)** The U87-MG spheroids stained with Hoechst (blue-all cells), Calcein-AM (green-live cells), Propidium iodide (red-dead cells), scale bar 200 µm. NMC and DMS represent no media change and daily media change respectively. **(C)** Size distribution of U87-MG spheroids (n>90) of the flow device (blue) and static device (green) on day 7.

We observed no change in well occupancy in our perfusion device indicating that the spheroids were not displaced after introducing the flow (data not shown). Live/dead assay (Hoechst, Calcein AM, and PI staining) performed on day 8 confirmed the high viability and low cell death in the spheroids cultured in flow devices, similar to that of the static devices with regular media change (Figure 4B). High fluorescence in Calcein-channel (green) and low fluorescence in PI-channel (red) clearly established that for both U87-MG and hMSCs (Supplementary Figure 2E), the cells in the spheroids formed in the device with continuous perfusion were alive and the dead cell population was low. Whereas when the culture media was not changed for 2 and 4 days consecutively, it resulted in high cell death compared to flow and static conditions where the culture media was replenished over time. This demonstrates that the perfusion system supports long-term spheroid culture with minimal user intervention. It is also interesting to note the formation of necrotic core in U87-MG tumoroids (Figure 4B) but not in the hMSCs spheroids (Supplementary Figure 2E). The average spheroid diameter was found to be 380 ± 16 µm and 154 ± 19 µm for U87-MG and hMSC spheroids respectively (Figure 4C, Supplementary Figure 2F) which might be the reason of this observation. We found that the average diameter of the flow device was comparable to the static device for both U87-MG and hMSC spheroids (Figure 4C, Supplementary Figure 2F). This implies that the flow of media did not interfere with the spheroid growth while maintaining the culture.

### 5. Anti-cancer drug testing on tumoroids

The increasing interest in 3D spheroid culture can be attributed to their suitability for applications in drug testing and cytotoxicity assays [49] [50]. Studies have demonstrated that 3D spheroids exhibit differential responses to specific drugs compared to 2D monolayer cultures, primarily due to variations in drug penetration and cell-cell interactions [51] [52]. Hence, to validate our device, we evaluated the effects of Temozolomide (TMZ), a widely used chemotherapeutic agent for glioblastoma (GBM), on U87-MG glioblastoma cells cultured as spheroids using S2 devices, using ultra-low attachment (ULA) plates, and as 2D monolayer.

To determine the IC_50_ values of anticancer drugs and to compare among various experimental conditions, U87-MG glioblastoma spheroids were cultured for 7 days using the S2 device and ULA plates, followed by a 48 h treatment with Temozolomide (TMZ) (Figure 5A). In parallel, U87-MG cells were cultured in a 2D monolayer using 96-well plates to compare drug toxicity between 2D and 3D cultures. After 48 hours of drug treatment the MTT assay was done and, the IC_50_ value was determined by fitting the datapoints in three parameter logistic regression model. 3D spheroid cultures exhibited significantly higher IC_50_ values compared to 2D monolayers (approximately 379µM, hill coefficient 1.17) (Figure 5B), consistent with previous findings [53] [54]. The IC_50_ value for spheroids generated in the S2 device and ULA-cultured spheroids was found to be approximately 684 µM (Hill coefficient 1.32) and 586 µM (Hill coefficient 0.65) respectively (Figure 5B-C). This difference might be attributed to the size difference of the spheroids formed in the S2 device and ULA plates. The S2-formed spheroids were smaller and compact compared to the spheroids formed in ULA plates, shown in Figure 3E, G. Furthermore, these results confirm that the quality of spheroids formed in the device is comparable to those generated in conventional ULA plates, highlighting its suitability for drug testing applications and supporting its potential as an alternative to ULA-based methods.

**Figure 5:**
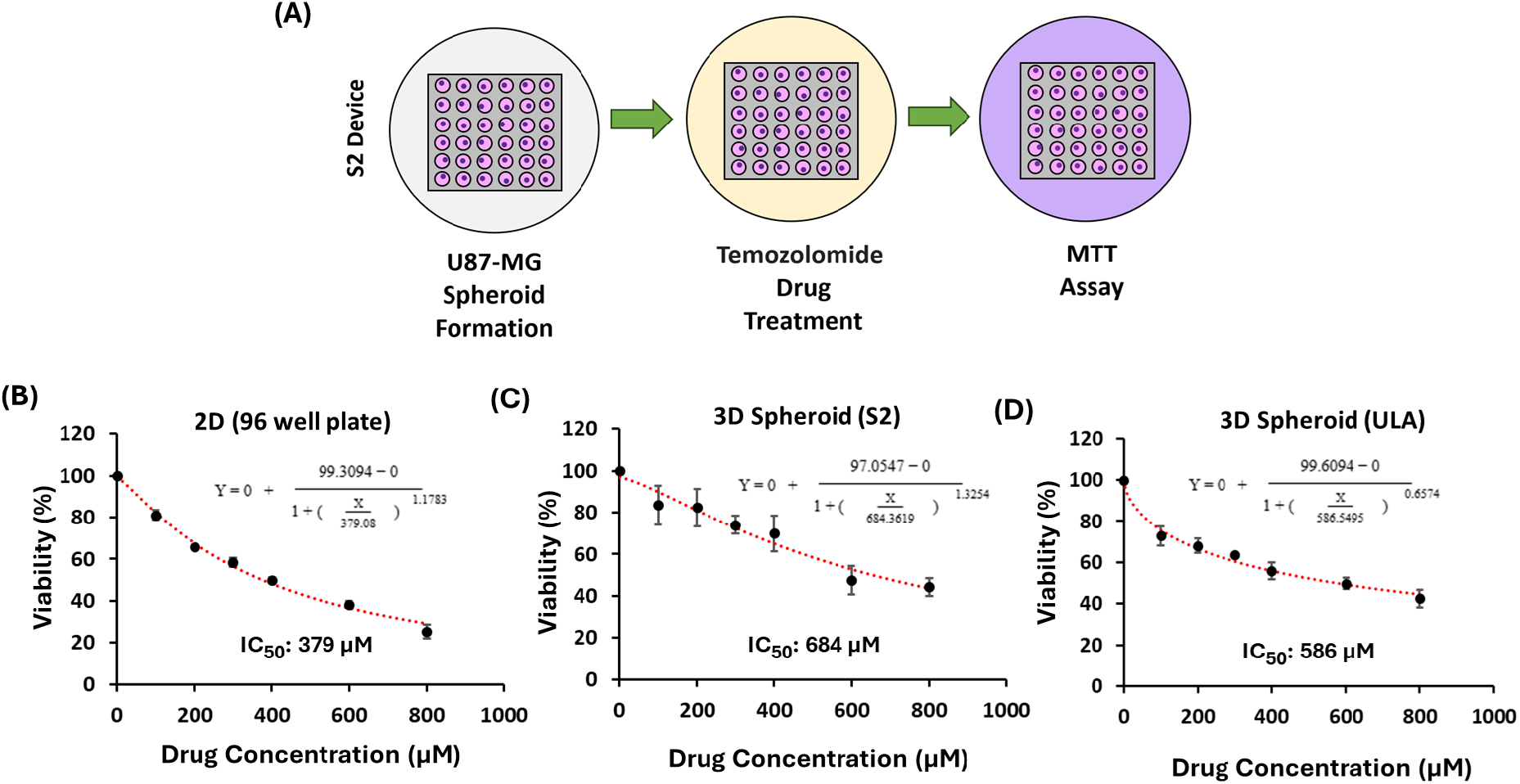
Validating the Device for Drug Testing: **(A)** Schematic of spheroid culture, drug treatment and MTT assay. **(B-D)** The percentage viability of U87-MG spheroid with varying concentration of TMZ (up to 800 µM) for (B) 2D-monolayer, (C) S2 devices, and (D), and in ULA plate has been shown. Data represents mean ± standard deviation (SD) from three technical replicates.

### 6. Gradient-sized spheroid formation in the S1 agarose device

Different studies have demonstrated that drug action, penetration, and cellular activity strongly depend on spheroid size. Larger spheroids impose greater resistance to mass transport, exposing cells to concentration gradients of nutrients, pH, and oxygen, which result in pronounced differences in biological responses. Gong et al. have shown the differential penetration of Doxorubicin (Dox) in different-sized spheroids composed of 8000 and 2000 cells. After 4 h of treatment the confocal imaging showed lesser Dox penetration in larger sized spheroids where the Dox signal was restricted to 100 µm from the boundary. Whereas, in smaller-sized spheroids, the penetration almost covered the entire spheroid [55]. In another work, Rovere et al., have shown that large size spheroids of mesenchymal stem cell (MSCs) show higher senescence, enhance the release of pro-inflammatory and pro angio-genic factors compared to their small sized counterparts [43]. Hence, generating spheroids of different but well-controlled size distribution is critical. Producing gradient-sized spheroids using conventional methods, such as the hanging drop technique, ultra-low attachment (ULA) plates, micropatterned plates, and liquid overlay methods, is challenging and requires labour-intensive adjustments of cell numbers and volumes for each desired size. Although microfluidic devices enable the production of spheroids with controlled sizes, they often necessitate complex dilutions, precise flow control, and expert handling, which limit their throughput and accessibility [27]. In contrast, the present device allows generation of gradient-sized spheroids without needing any additional instrumentation.

To generate the gradient sized spheroids, we used an alternative material, agarose which is easy to mold, mechanically stable enough to preserve intricate structures, and inherently non-adhesive to cells, making it suitable for culturing spheroids [29] (see Materials and Methods for details). For generation size-gradient spheroids, we tilted the devices at different angles instead of keeping them flat followed by seeding of HTR-8 cells. This tilt created a gradient in media height across the rows of the device. As a result, more cells settled in the bottom rows where the media height was greater, while fewer cells reached the top rows due to the lower media level (Figure 6A). With careful and methodical optimization (data not shown), an inclination of 20^0^ gave rise to a consistent gradient in spheroid sizes, gradually increasing from row 1 to row 11. As one may expect intuitively, the average diameter of the spheroids in row 6 shows similar values for all the inclinations, which is similar to the mean diameter of the flat device, indicating that the media column height for the middle row is identical for all the cases shown.

**Figure 6:**
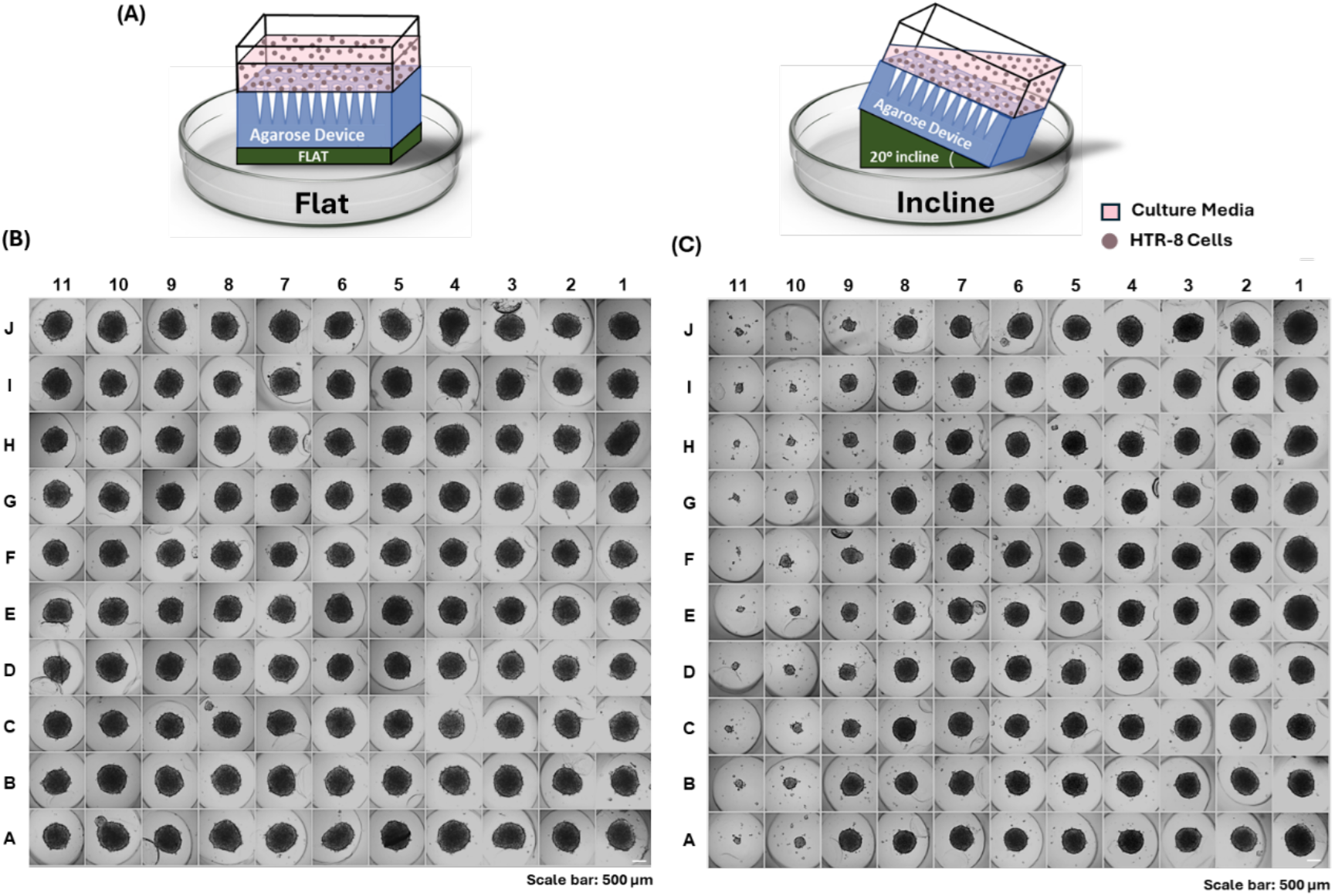
Gradient-sized spheroid formation in S1 devices: (A) Schematic representation of the inclined setup for flat (0°), and 20° inclinations. Representative phase contrast images (from three independent experiments) of HTR-8 spheroids on Day 4 (B) Flat (left) vs (C) Inclined at 20° (right) has been shown. Scale bar: 500 µm.

### 7. 3D Co-culture of placental trophoblasts in S1 device

While spheroid size is crucial, co-culturing multiple cell types in 3D spheroids further improves the ability to mimic disease environments. Co-culture systems are being used to model breast cancer [56], pancreatic cancer [57], bacterial infection in tumoroids [58], etc. For example, to study various disease conditions during pregnancy, it is important to mimic the placental physiology. Traditional 2D co-cultures of trophoblasts provide insights into cell–cell interactions but lack the complex architecture of the maternal–fetal interface. In contrast, 3D co-cultures overcome these limitations by recreating spatial gradients, cell–matrix interactions, and physiological diffusion zones [59]. Several studies have reported 3D trophoblast spheroid or organoid models that form invasive and hormone-secreting structures [60] [61] [62].

To establish the co-culture in our device, two types of trophoblast cells, extravillous trophoblasts (HTR-8) and choriocarcinoma derived trophoblasts (BeWo) were sequentially introduced in the device. For sequential seeding, one cell type was introduced in flat condition, resulting in uniform seeding, and the other cell type in gradient manner, resulting into formation of size gradient. HTR-8 (labelled with Green Cell tracker) formed the core while BeWo (labelled with Red Cell tracker) occupied the periphery of the co-cultured spheroid. To evaluate spheroid architecture and cellular interactions confocal images were taken at day 1 (24 hours post BeWo seeding) and at day 5.

For the first condition, HTR-8 cells (green) were seeded and allowed to form spheroids till day 4 in gradient condition. On day 4, BeWo (red) were seeded in flat condition. This was considered as day 1 of co-culture. The two cell types in this model were seen as distinct layers (shown in Figure 7A-i). BeWo cells (red) were observed covering the periphery of the pre-formed HTR-8 spheroids (green). This observation indicated initial localization without substantial mixing with the existing HTR-8 spheroid (shown in Figure 7A-i). At day 5, the co-cultured spheroids showed an integration between the two cell types. The merged z-stack projections in Figure 7A-ii exhibit a substantial overlap and intermixing at the HTR8-BeWo interface which suggests an active cellular interaction and possibly migration across the boundaries.

**Figure 7:**
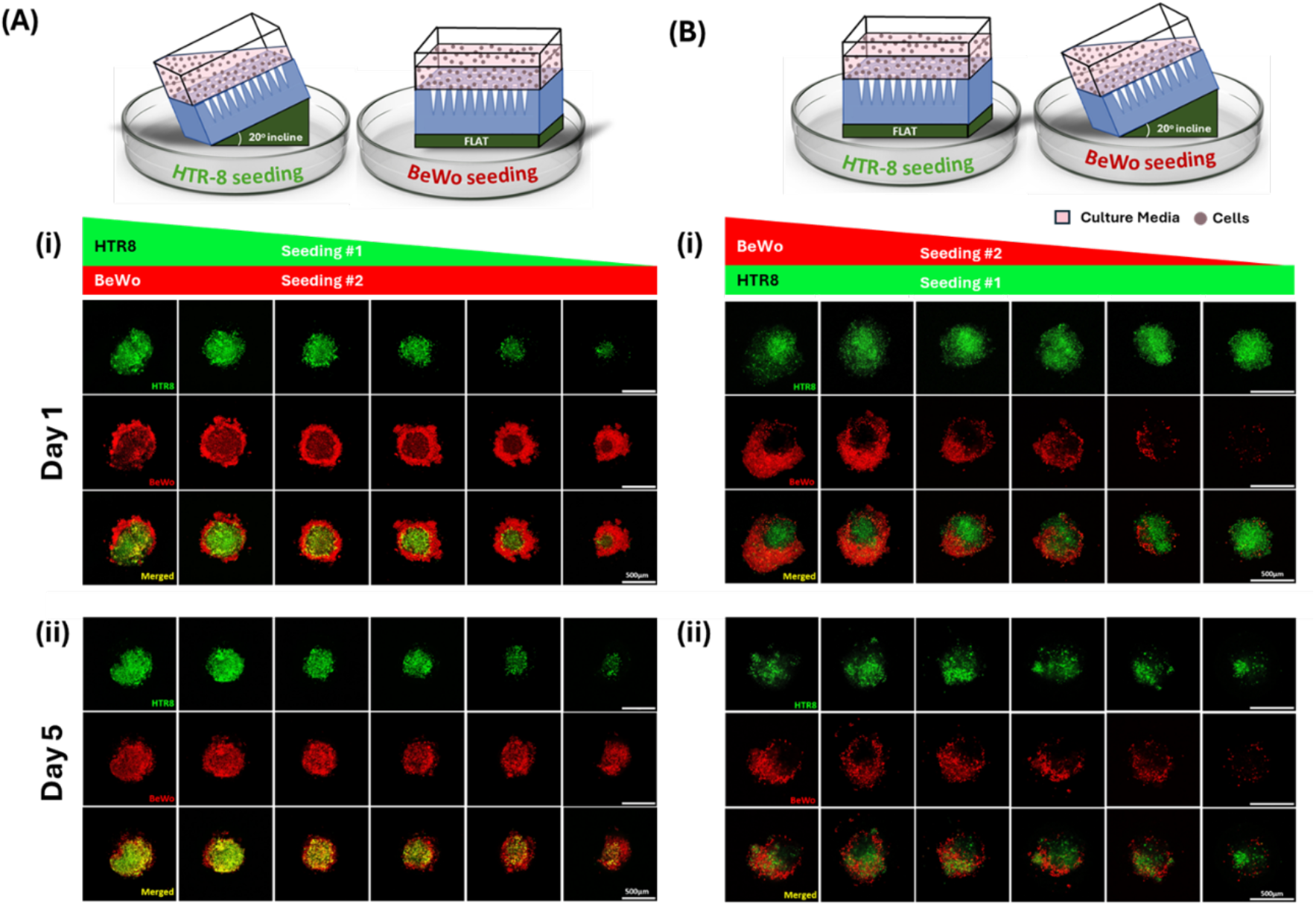
3D spheroid co-culture of placental trophoblasts: HTR-8 and BeWo cells were seeded sequentially in the S1 spheroid devices. The maximum intensity projection of the confocal images of the co-cultured spheroids are represented from three technical replicates. (A) Fluorescence images of the HTR-8 spheroids (green), seeded first in gradient condition followed by BeWo (red), seeded in the flat condition are shown. Images were taken at (i) day 1 and (ii) day 5 post seeding of BeWo cells. (B) Fluorescence images of the HTR-8 spheroids (green), seeded first in flat condition followed by BeWo (red), seeded in the gradient condition are shown. Images were taken at (i) day 1 and (ii) day 5 post seeding of BeWo cells. Scale bar: 500 µm.

For the second condition, HTR-8 cells (green) were seeded and allowed to form spheroids till day 4 in flat condition. On day 4, BeWo (red) were seeded in gradient condition. This time at day 1, BeWo cells were found to be deposited on one side of the HTR-8 spheroids (Figure 7B-i) unlike the first condition (Figure 7A-i) where it covered the existing spheroids from all sides. This deposition might be a result of the incline seeding technique. By day 5, the both BeWo and HTR-8 were intermixed with each other (Figure 6B-ii) similar to as observed in the first condition (Figure 6A-ii).

In both the cases, the compaction and cell intermixing observed by Day 5 align with known placental physiology, where extravillous trophoblasts and syncytiotrophoblasts interact during villi formation and maturation in the process of placentation [63]. It is also observed that the HTR-8 cells being migratory in nature are seen to disperse and penetrate through the BeWo layer. This model can further be used to study pregnancy related disorders like implantation failure, preeclampsia, and placental toxicity. Although trophoblast cells were used to model placenta-relevant physiology, the results demonstrate the device’s ability to support 3D co-culture conditions and suggest its wide applicability in studying cell–cell interactions in other contexts such as cancer, infections, and more.

## Critical Evaluation and Conclusion

| Platform | Cost (USD) | No. of spheroids | Cost/Spheroid (USD) |
| --- | --- | --- | --- |
| **Ultra-low Attachment (ULA) Plates or Surface** |  |  |  |
| MicroTissues® 3D Petri Dish, 256 wells | 575 | 256 | 2.25 |
| 96-well Spheroid Microplates | 50 | 96 | 0.52 |
| Ultra Low Attachment surface, 4515, by Corning |  |  |  |
| AggreWell™800 6-well plate starter kit | 243 | 18000 | 0.014 |
| **Hanging Drop Plates** |  |  |  |
| Perfecta3D™ Hanging Drop Plate | 26 | 96 | 0.27 |
| **S1 Device** | 1.16 | 100 | 0.0116 |

The high-throughput 3D spheroid generation device presented in this study offers numerous advantages but also has certain limitations. A primary concern is the potential for drug adsorption on PDMS surfaces and absorption into agarose, particularly for hydrophobic molecules, which may affect the accurate determination of free drug concentrations. Additionally, when drug testing is performed directly within the device, all 100 spheroids share the same media. While this uniformity strengthens statistical analyses, it limits the number of experimental conditions that can be assessed simultaneously. However, as discussed, the spheroids can be easily harvested and redistributed into conventional multiwell plates for further testing, addressing this limitation.

Despite these challenges, the device offers significant advantages over existing methods for spheroid generation. It is cost-effective, easy to fabricate, and scalability, producing uniformly sized spheroids without the need for complex equipment or specialized expertise. The device enables the generation of hundreds of spheroids from diverse cell types, including primary and cancer cell lines, and supports long-term culture with minimal intervention through an integrated perfusion system. Its utility as a drug-testing platform was validated using temozolomide on U87-MG cells, yielding IC_50_ values consistent with those obtained using standard methods such as ultra-low attachment (ULA) plates. Furthermore, the device can be easily modified to produce size-gradient spheroids, and has can provide a simple and efficient platform for 3D co-cultures to model cell-cell interactions with spatial control over cell density.

In conclusion, the high-throughput, low-cost, DIY device described in this paper addresses key limitations of existing spheroid generation techniques. We believe it will be particularly beneficial for resource-limited laboratories while also offering innovative solutions to technical challenges faced by advanced high-throughput systems.

## Acknowledgments

We would like to thank Dr. Deepak Modi, his student Mr. Anshul Bhide, NIRRCH-ICMR, and Prof. Sudhir H Ranganath (Siddaganga Institute of Technology, Tumkur, India) for evaluating the devices independently which helped to increase our confidence on the usefulness of the device. The placental cell lines were generously provided by Dr. Modi. The U87-MG cell lines were generously provided by Dr. Shilpee Dutt (JNU, India). AM acknowledge funding support from IMPRINT (IMP/2019/000115), WRCB (DO/2021-WRCB002-066), ICMR (17X(3)/Adhoc/65/2022-ITR). SM, DR, and PM thank ICMR (17X(3)/Adhoc/65/2022-ITR), WRCB (DO/2021-WRCB002-066), and IRCC (IIT Bombay) for providing fellowship support, respectively. All the authors thank IIT Bombay for providing infrastructural support.

## Authors Contribution

PM, SM, DR, AM conceptualized the project. PM, SM, TG, DR, AV, SK, developed the methodology. PM, SM, TG, DR, AV, SK, VS performed the experiments, collected and analyzed the data. SM, DR, AV, TG, PM, and AM wrote and reviewed the manuscript. AM managed the funding, resource allocation, and overall supervision.

The authors declare no conflict of interest.

