## Supplementary Fig. 1 for "Shape, Shrink, Spheroid: A DIY High-throughput Spheroid Generation Device"

### Supplementary Figure 1:

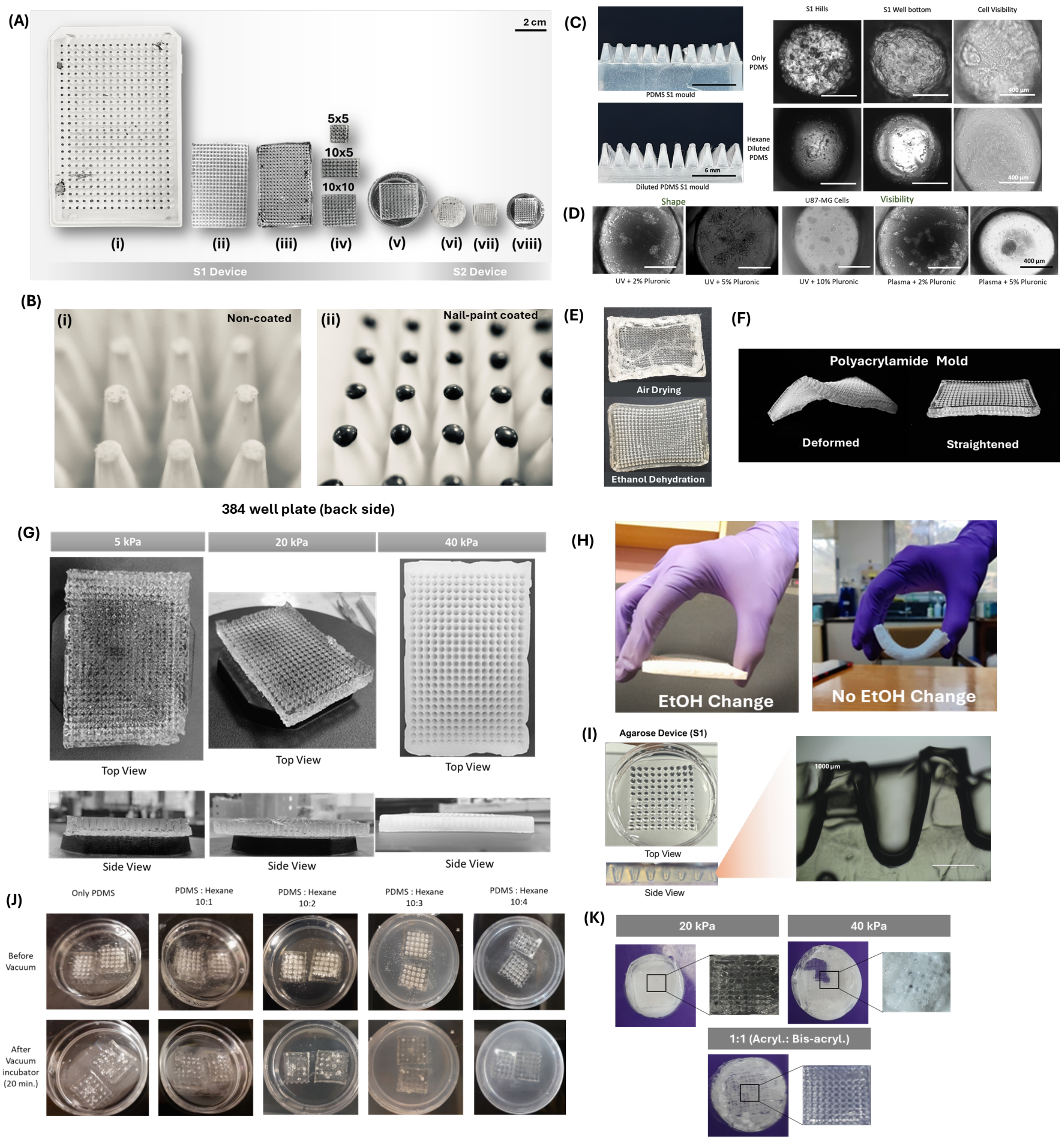


**Supplementary Figure 1: (A)** The actual images of the fabrication steps have been represented, (i) back side of a nail-paint coated 384 well plate, (ii) shrunk PA mold with 384 wells (before straightening, see the text for more details), (iii) PDMS mold with 384 hills, (iv) PDMS hills has been cut to various shapes such as 5x5, 10x5, 10x10 etc., for fabrication of device with any AxB shape, (v) 10x10 PDMS device (S1), (vi) PA mold for S2 device, (vii) PDMS mold with hills for S2 device, (viii) 10x10 S2 PDMS device. The images represented are of actual size with a scale bar of 2cm. **(B)** i. Flat apex of 384-well plate bottom structures without any coating, ii. Round apex of 384-well plate bottom structures after nail-paint coating, **(C)** The difference between using undiluted and hexane-diluted PDMS while fabricating the PDMS mold (with hills) is shown. The well apex (scale bar 6 mm), microscopic images of well shape (scale bar 400 µm) and visibility are shown. **(D)** The U87-MG cell attachment on day 4 with Pluronic coating upon varying concentration in presence of UV and plasma treatment has been shown, scale bar 400 µm. **(E)** The PA mold after air drying and ethanol drying has been shown. **(F)** The deformed shrunk PA mold before was straightened by keeping it in a moist chamber. **(G)** The PA gel compositions with different stiffness are represented, **(H)** The importance of EtOH change has been shown. The resultant rigid PA mold after ethanol change and the soft PA mold with no EtOH change has been shown respectively, **(I)** Agarose device top view and side view have been shown along with a microscopic cross section of a single well has been shown, scale bar 1000 µm, **(J)** Images for PDMS dilution with various concentration of n-Hexane (i.e., 10:1, 10:2, 10:3, 10:4) has been shown **(K)** The images of 20kPa, 40kPa and 1:1 mixture of acrylamide (40%) and bis-acrylamide (2%) for the S2 mold generation has been shown.
