## Supplementary Fig. 2 for "Shape, Shrink, Spheroid: A DIY High-throughput Spheroid Generation Device"

### Supplementary Figure 2:

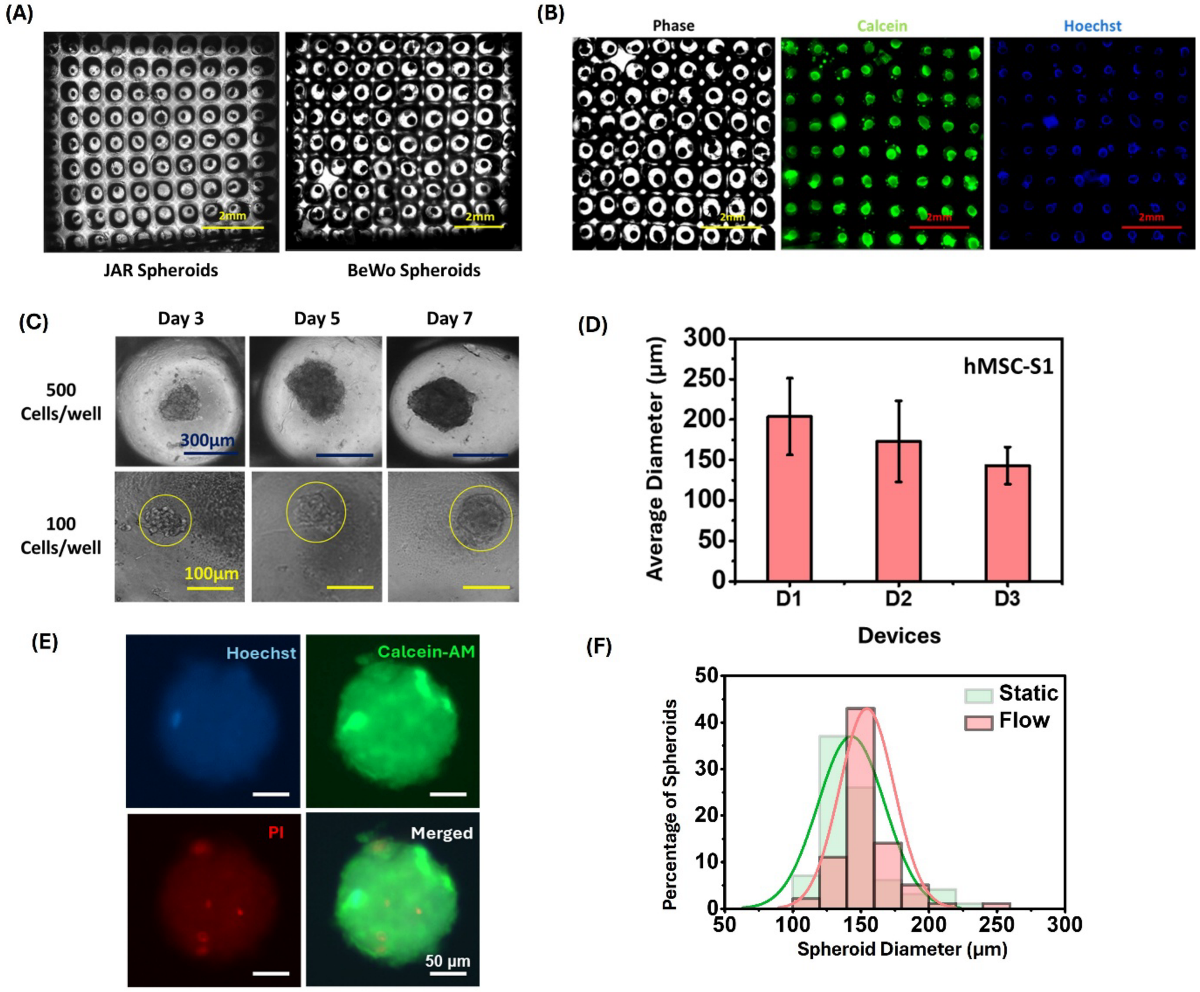


**Supplementary Figure 2: (A)** Scanned images of the S1 device with placental cell lines JAR and BeWo are shown, indicating the spheroid forming capability of the device. **(B)** The imageability of the device has been shown with scanned images of day-7 U87-MG spheroids in phase contrast and stained with Calcein (green) and Hoechst (blue), scale bar 2 mm. (C) The spheroid forming capability with low cell number such as 500 and 100 cells/well is shown at day 7, scale bar 300 µm and 100 µm respectively, (D) Device to device variation in hMSC spheroid diameter has been shown, D1, D2 and D3 denotes Device 1,2 and 3 respectively, (E) Fluorescent hMSC spheroids of day 7 in flow device has been shown. The spheroids were stained with Calcein (green), Hoechst (blue) and PI (red), scale bar 50 µm, (F) The distribution of hMSC spheroid diameter on Day 7 in flow device (red) has been shown and compared with S1 static device (green) by overlaying in a single graph.
